# Overcoming the Woody Barrier: Dodder Enables Efficient Transfer of Infectious Clones to Woody Plants

**DOI:** 10.1101/2025.01.14.633052

**Authors:** Andrea Sierra-Mejia, Mohammad Hajizadeh, Habeeb Yinka Atanda, Ioannis E. Tzanetakis

## Abstract

Woody hosts are notoriously resistant to genetic transformation. Traditional methods, such *as Agrobacterium-*mediated transformation, are often inefficient, and this limitation extends to delivering infectious clones to woody plants. Dodder species (*Cuscuta spp.)* are holoparasitic plants that can establish direct connections with the vascular tissue of the parasitized plants, allowing them to facilitate virus transmission between unrelated botanical species. We demonstrated that a novel dodder-based approach achieved superior transmission in *Rubus* spp. compared to direct agroinoculation. The transmission rates for systemic blackberry chlorotic ringspot virus transmission increased from 9% to 73%, whereas the transmission of the phloem-restricted blackberry yellow vein associated virus rose from 0% to 46%. This novel method expands the toolbox available to plant biologists to study virus-host interactions in woody plants.

## 1. Introduction

Infectious clones are essential tools for studying viruses and virus-host/vector interactions, as they offer a fully characterized, single-virus source that can be modified to appropriately address the posed hypotheses (Abrahamian et al., 2020; Pasin et al., 2019). Infectious clones are delivered to host cells by a variety of methods but primarily rely on transient plant transformation techniques. These methods include biolistic bombardment, protoplast transformation, and *Agrobacterium*-mediated transformation, hereafter referred to as agroinoculation, (Abrahamian et al., 2020; Canto, 2016; Chincinska, 2021; Nagyova & Subr, 2007), with the latter typically recognized as the most effective approach (Altpeter et al., 2016).

While many infectious clones have been successfully introduced to herbaceous hosts, delivering them to woody hosts poses a significant challenge (Abrahamian et al., 2020; Dawson & Folimonova, 2013; Pasin et al., 2019). Woody plants are recalcitrant to genetic modification (Dawson & Folimonova, 2013), often resulting in inefficient agroinoculation (Abrahamian et al., 2020; Zhang & Jelkmann, 2017). Moreover, the phloem-restricted nature of certain viruses presents greater challenges in woody hosts compared to herbaceous plants (Orílio et al., 2014; Qiao & Falk, 2018; Shi et al., 2016; Wang et al., 2009). In general terms, direct agroinoculation is inefficient in woody hosts, often requiring modifications of protocols, expertise in plant *in vitro* techniques, and specialized equipment. As a result, this approach is typically only available for a limited number of crops (Cui et al., 2018; Muruganantham et al., 2009; Kurth et al., 2012; Zhang & Jelkmann, 2017). For example, infectious clones of citrus yellow vein clearing virus and apple chlorotic leaf spot virus have only been successfully delivered to citrus and apple seedlings, respectively, using a modified vacuum infiltration protocol (Cui et al., 2018; Zhang & Jelkmann, 2017). In contrast, particle bombardment has been used to deliver infectious clones of prunus necrotic ringspot virus and plum pox virus to peach seedlings (Cui & Wang, 2017). Delivery of infectious clones to grapevine is even more challenging. To achieve infections with grapevine leafroll-associated virus-2 and grapevine virus A, scientists have adapted vacuum infiltration and agro-drenching protocols and utilized micropropagated seedlings (Kurth et al., 2012; Muruganantham et al., 2009). Nevertheless, there are straightforward success stories, such as the direct leaf agroinfiltration of citrus leaf blotch virus in citrus trees (Vives et al., 2008); however, these cases remain exceptions rather than the norm. Current protocols pose limitations when it comes to woody plants, emphasizing the need for a universal, efficient, and user-friendly method for delivering infectious clones to recalcitrant hosts.

The ability of dodder (*Cuscuta spp.*) to bridge virus transmission amongst unrelated botanical plant species was first described in 1940 (Bennett, 1956). Dodder is a holoparasitic plant that relies exclusively on its host for nutrient acquisition. It grows as tendrils that wrap around and climb their host plants. They use a specialized haustorium structure to penetrate the host’s tissues and acquire nutrients. The haustorium penetrates the host tissue forming a direct connection with the vascular tissue of the parasitized plant. This interaction creates a bidirectional pathway that allows for the exchange of proteins and RNA, effectively acting as a bridge between dodder and its host (Bhat & Rao, 2020; Hosford, 1967; Li et al., 2022). Dodder can attach to multiple host species simultaneously, facilitating the transmission of pathogens from infected to healthy plants. This transmission can occur even between unrelated plant species, where grafting typically would not facilitate pathogen movement (Li et al., 2022, Mikona & Jelkmann, 2010). Using dodder to facilitate transmission is particularly effective for viruses with unknown vectors or those that cannot be mechanically inoculated, such as phloem-restricted viruses (Bennett, 1956; Mikona & Jelkmann, 2010).

Here, we present a dodder-based approach for efficiently transferring infectious clones to woody hosts. Infectious clones are delivered to an herbaceous host via leaf agroinoculation, and then dodder is employed to transfer the clones to the target host. We demonstrated the species-independent nature of the approach using infectious clones from viruses belonging to two genera, one systemic and one phloem-restricted, across two different plant genera: *Nicotiana benthamiana* and *Rubus* spp.

## 2. Materials and methods

### 2.1 Agroinoculation

As a baseline, we directly delivered infectious clones of blackberry chlorotic ringspot virus (BCRV) and blackberry yellow vein associated virus (BYVaV) (Sierra-Mejia, 2025a;b) to their woody host: *Rubus spp*. Three distinct approaches—agro-paste, syringe, and vacuum inoculation—were utilized. The infectious clones were prepared using equal volumes of their genomic RNAs and *Agrobacterium* at an optical density of OD600 = 1.0. The agro-paste approach was conducted as described by Villamor et al (2021), whereas the syringe inoculation followed the protocol outlined by Sierra-Mejia et al (2025a). In vacuum inoculation, the inoculum was prepared using the syringe technique and applied to the leaves of young seedlings. These seedlings were immersed in the inoculum within a Scienceware vacuum desiccator connected to a GAST vacuum pump, where they were subjected to a pressure of 600 mmHg for 5 minutes. The seedlings were then transplanted into soil.

For the dodder approach the BCRV and BYVaV infectious clones (PJL89-BCRV and PJL89-BYVaV) were delivered to *Nicotiana benthamiana* as described Sierra-Mejia et al (2025a;b). After inoculation, plants were maintained in growth chambers under a 16/8h light/dark photoperiod at 22 °C. Systemic virus infection was assessed through RT-qPCR for BCRV and BYVaV, following the protocols outlined by Sierra-Mejia et al. (2025a;b) (**Supplementary Table 1**).

### 2.2 Dodder transmission

For the dodder-inoculation, *Cuscuta pentagona* seeds were scarified for 20 min in 95% sulfuric acid, followed by three 10-minute water washes. Seeds were placed in soil (less than 0.5 cm deep) next to 4-to 6-week-old indicator plants, with germination observed within a week. Dodder was allowed to colonize the indicator plants for 3- to 4-weeks. This first step ensures the establishment and maintenance of virus-free dodder for subsequent transmission experiments.

Next, syringe-inoculated plants were placed next to a dodder-infested indicator and dodder was allowed to establish on the virus-positive plants. Once the dodder had successfully taken over the herbaceous plant, the targeted hosts—*Rubus spp*. was introduced, allowing the tendrils to move and establish onto the recipient host **(Figure 1**). Additionally, for the phloem-restricted BYVaV, we introduced *N. benthamiana* to assess the transmission efficiency and to determine whether it improved compared to the standard method (**Figure 1**).

**Figure 1.**
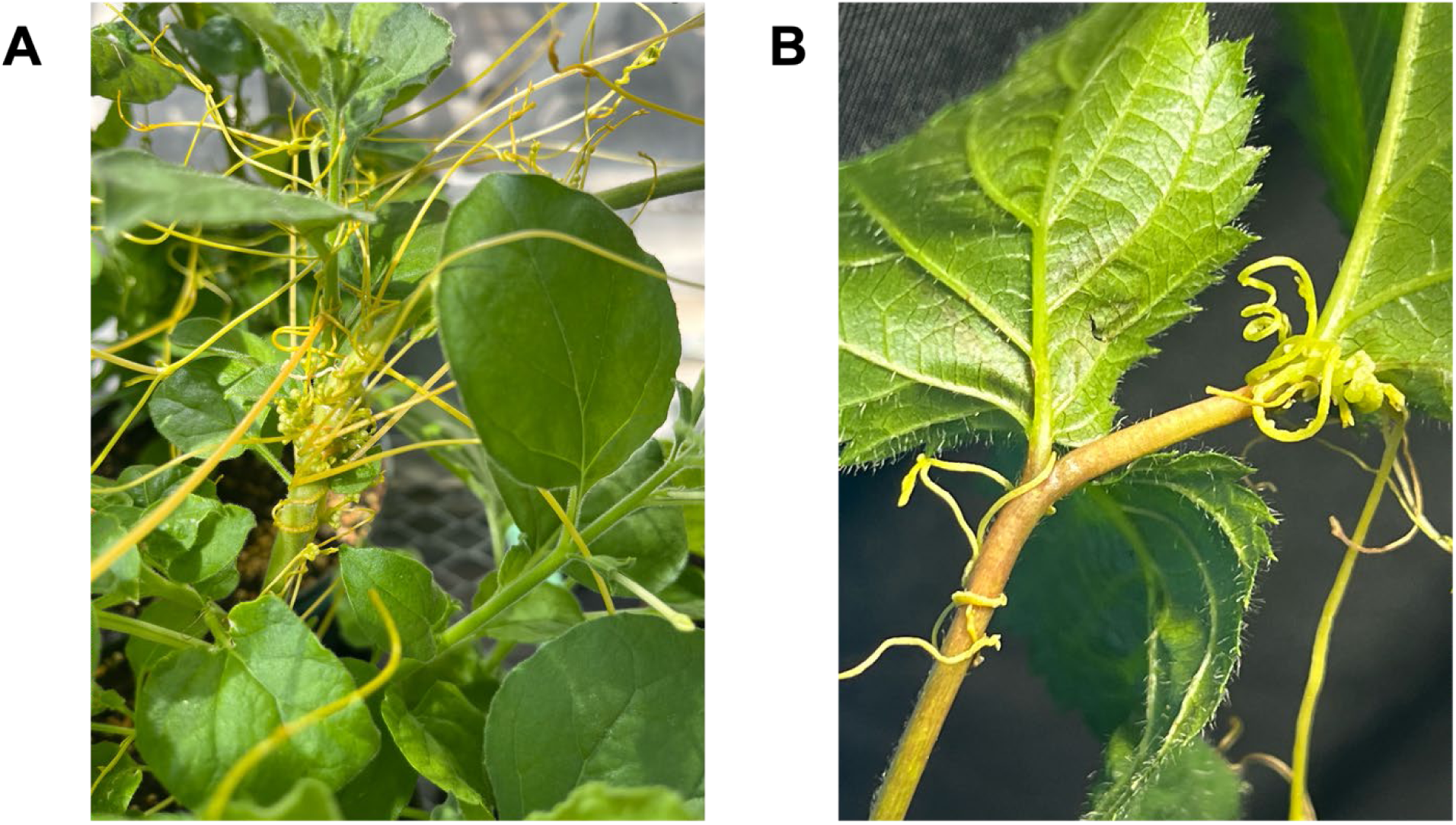
Dodder (*Cuscuta pentagona*) attachment to two distinct hosts. A) *Nicotiana benthamiana* and B) *Rubus* spp.

Dodder was detached from the recipient plants after initial establishment: one month for BCRV and three months for BYVaV, as the latter requires more time to establish due to its phloem-restricted nature. All plants were kept in an insect-proof greenhouse, which allowed for the provision of natural lighting conditions (Anonymous, 2019). Temperatures were kept between 20 and 30°C, and relative humidity was maintained around 60% to provide optimal conditions for dodder growth (Mikona & Jelkmann, 2010).

### 2.3 Detection

Virus replication is reduced at higher temperatures (Constable et al., 2013; Glasa et al., 2003; Hily et al., 2016) and for this reason *Rubus* plants were placed in a cold room at 4 °C and kept in the dark for about one month to simulate chilling. After this period, the plants were moved to growth chambers, where they were maintained under a 16/8 h light/dark photoperiod at 22 °C. This step was designed to promote systemic virus movement and increase viral titer. Transmission was then assessed using RT-qPCR, as previously described (Sierra-Mejia et al., 2025a;b). RT-qPCR results were further validated using RT-PCR, as previously described, followed by Sanger sequencing (Poudel et al., 2013; 2014).

### 2.4 Statistical analysis

The qPCR results were compiled into a dataset that categorized samples by transmission method and the presence or absence of successful transmission. A contingency table was created to summarize the data, and Fisher’s Exact Test was conducted to assess whether a significant association existed between the transmission method and transmission outcome. A one-tailed p-value was then calculated based on the results of Fisher’s test (P < 0.05). All analyses were performed using R version 4.4.1(R Core Team, 2023).

## 3. Results

### 3.1 Direct agroinoculation

Direct agroinoculation of BCRV into *Rubus spp.* was successful only with the agro-paste method, resulting in virus establishment in 3/33 plants (9%). In contrast, neither the syringe method nor the vacuum inoculation resulted in infection (0/12 plants of each treatment), whereas herbaceous indicators consistently tested 100% positive after syringe inoculation (Sierra-Mejia et al., 2025a). For BYVaV, none of the employed methods were successful for *Rubus* (0/21 plants), whereas inoculation onto *N. benthamiana* yielded a success rate of 56% (10/18 plants; Sierra-Mejia et al., 2025b) (**Table 1**).

**Table 1.**
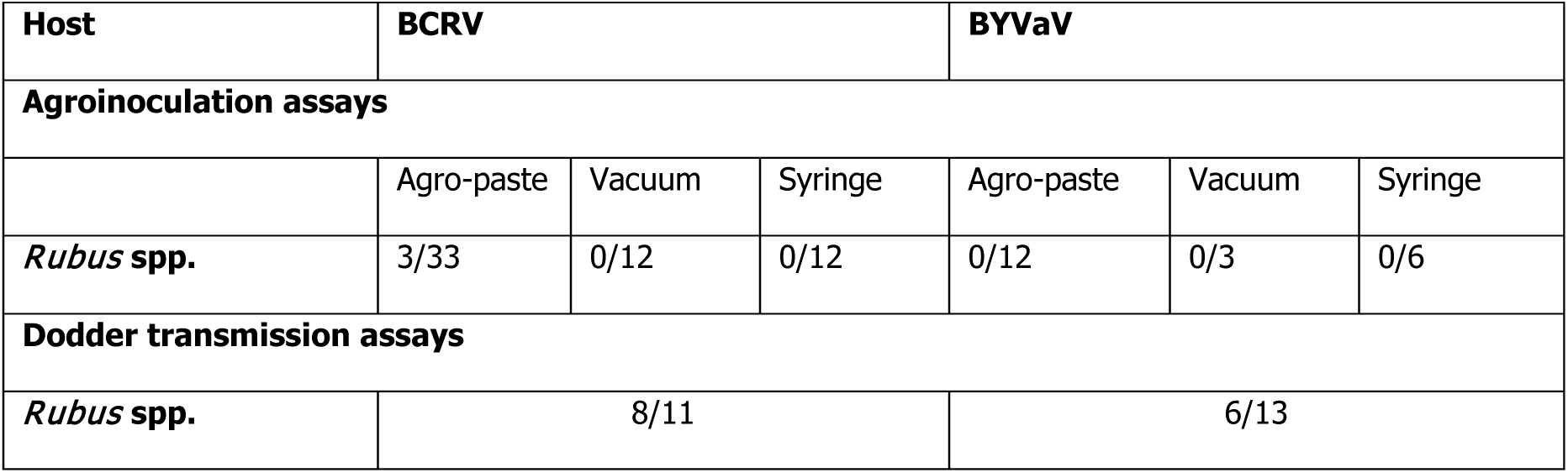
Blackberry chlorotic ringspot virus (BCRV), and blackberry yellow vein associated virus (BYVaV) detection in agro-inoculated and dodder transmission assays on woody hosts.

| Host | BCRV |  |  | BYVaV |  |  |
| --- | --- | --- | --- | --- | --- | --- |
| Agroinoculation assays |  |  |  |  |  |  |
|  | Agro-paste | Vacuum | Syringe | Agro-paste | Vacuum | Syringe |
| *Rubus* spp. | 3/33 | 0/12 | 0/12 | 0/12 | 0/3 | 0/6 |
| Dodder transmission assays |  |  |  |  |  |  |
| *Rubus* spp. | 8/11 |  |  | 6/13 |  |  |

### 3.2 Dodder transmission

Among *Rubus spp.*, 8 out of 11 (73%) plants tested positive for BCRV, and 6 out of 13 (46%) tested positive for BYVaV. Additionally, 6 out of 10 (60%) *N. benthamiana* plants tested positive for BYVaV (**Table 1**). RT-qPCR results were verified using RT-PCR **(Figure 2**) and amplicons from both viruses were then Sanger sequenced, with all sequences being virus-specific.

**Figure 2.**
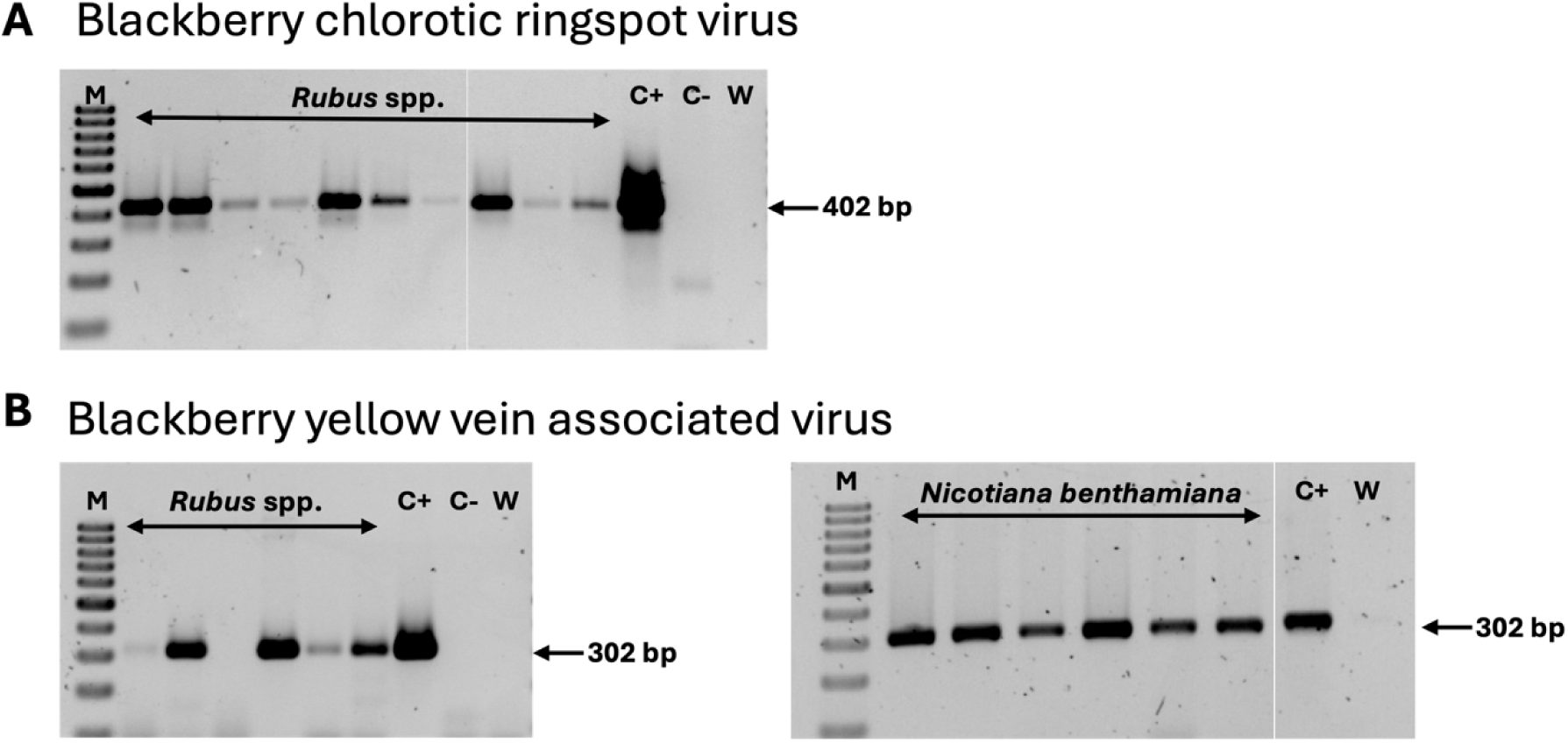
Detection of blackberry chlorotic ringspot virus (BCRV), and blackberry yellow vein associated virus (BYVaV), after dodder transmission assays using RT-PCR. A) BCRV and B) BYVaV. M: 100 bp molecular marker, C+: plant positive control, C-: plant negative control, and W: water control

### 3.3 Symptoms development

Similar to previous studies (Poudel et al., 2013; 2014), no symptoms were observed on *Rubus spp.* infected with BCRV and BYVaV. *N. benthamiana* plants infected with BYVaV via dodder exhibited symptoms like those seen in syringe-inoculated plants, including interveinal chlorosis, leaf yellowing, and mild curling **(Figure 3**).

**Figure 3.**
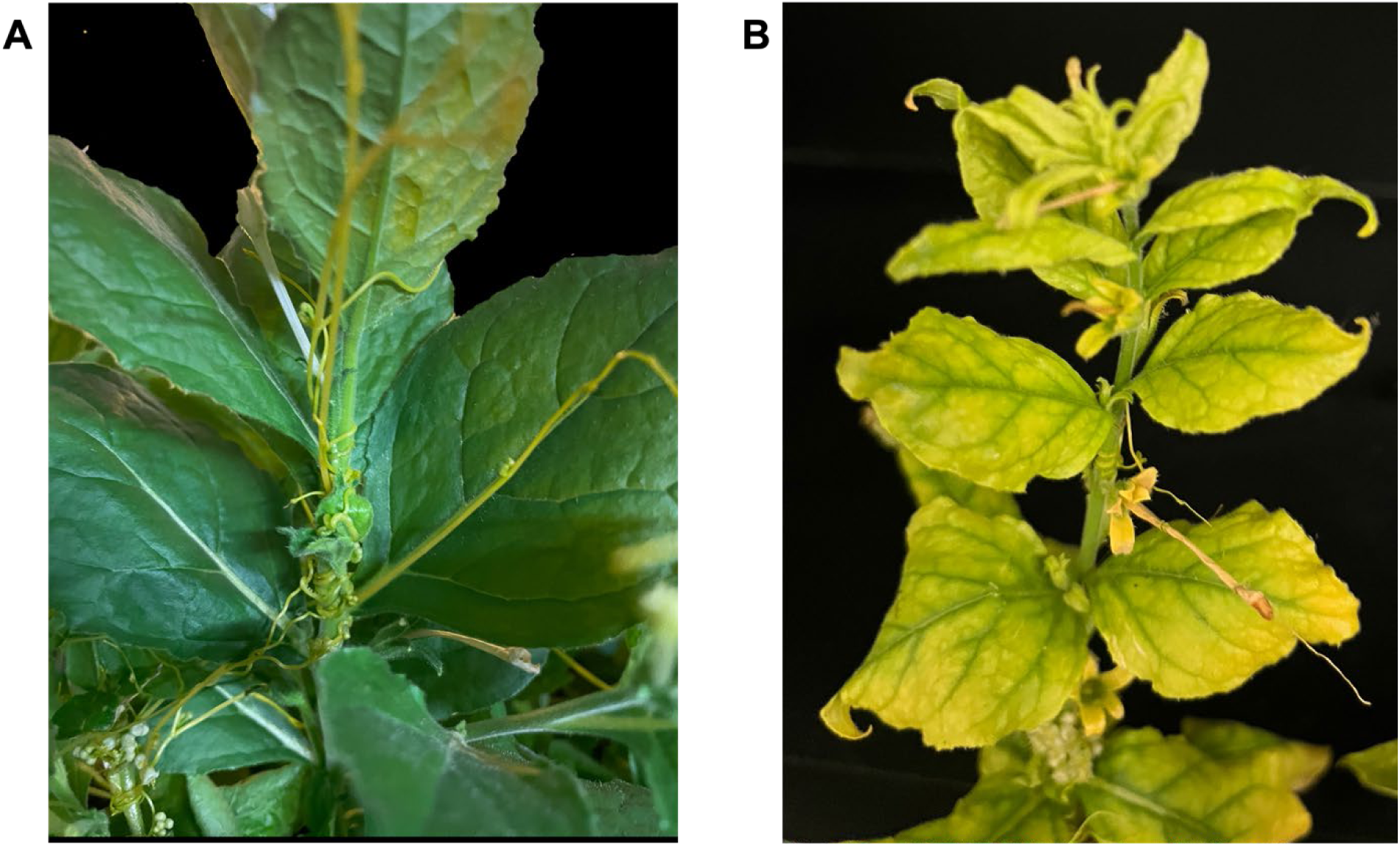
Symptomology of blackberry yellow vein associated virus (BYVaV) positive *Nicotiana benthamiana* after dodder transmissions assay. A) Healthy *N. benthamiana* with dodder attachment. B) BYVaV-positive *N. benthamiana* with dodder attachment exhibiting interveinal chlorosis, leaf yellowing, and mild curling.

### 3.4 Statistical analysis

For transmission experiments using *N. benthamiana*, the Fisher’s Exact Test revealed no statistically significant differences between the dodder method and the syringe agroinoculation method, with a one-tailed p-value of 0.5 (**Figure 4**).

**Figure 4.**
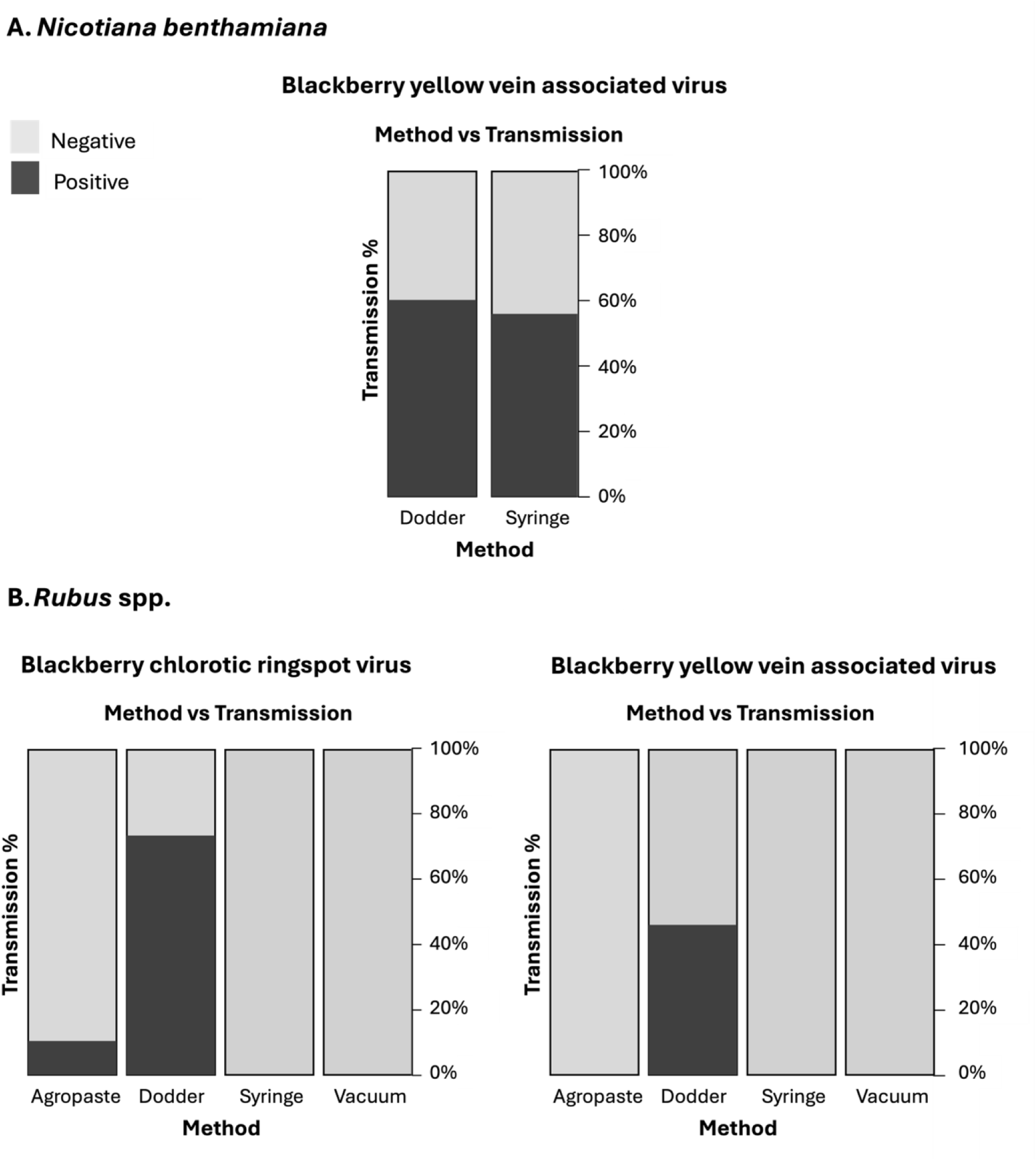
Plot illustrating the relationship between method and transmission. A) Transmission of blackberry yellow vein associated virus (BYVaV) in *Nicotiana benthamiana.* BYVaV transmission was observed with the syringe method at an efficiency of 56%, compared to 60% efficiency using the dodder method. B) Transmission of blackberry chlorotic ringspot virus (BCRV) and BYVaV in *Rubus* spp. Left, transmission of BCRV was observed with the agro-paste method at an efficiency of 9%, compared to 73% efficiency using the dodder method. Right, transmission of BYVaV was exclusively observed using the dodder approach, achieving an efficiency of 46%.

In contrast, transmission experiments using *Rubus* spp., revealed a statistically significant difference between the dodder and the agroinoculation methods for both BCRV and BYVaV transmission (**Figure 4**). The one-tailed p-value for BCRV transmission was 4.09e-06, whereas for BYVaV transmission, it was approximately 0.005.

## 4. Discussion

Direct agroinoculation of the woody host was successful, although with low efficiencies, for BCRV. However, agroinoculation of BYVaV was unsuccessful across all methods employed. This discrepancy could be due to the nature of tissue tropism between the two viruses: while both viruses are systemic, BCRV can move out of the phloem, whereas BYVaV is phloem-restricted. Methods such as syringe and vacuum inoculations can effectively deliver *Agrobacterum* to the mesophyll cells, but they encounter greater difficulty in reaching the phloem (Chincinska, 2021). Moreover, even in herbaceous hosts, BYVaV agroinoculation is less efficient compared to BCRV with BYVaV infectious clone showing 56% efficiency in *N. benthamiana* whereas the BCRV counterpart infected all *N. benthamiana* and *N. occidentalis* plants (Sierra-Mejia et al., 2025a;b).

Considering these results and aiming for a more efficient method to deliver infectious clones to woody plants, we evaluated a dodder-based transmission protocol. This approach utilizes dodder to facilitate virus transfer, thus simplifying existing delivery methods for woody plants. Factors such as *Agrobacterium* strain, plant genotype, leaf infiltrability, and developmental stage are critical for the success of agroinoculation (Chincinska, 2021; Karami, 2008; Salazar-González et al., 2023) but become unimportant for dodder attachment. This makes the described method superior when compared to previously described approaches.

As a proof of concept we explored the dodder-based method for facilitating infection of the more recalcitrant of the two viruses in the study, BYVaV, in *N. benthamiana.* Results indicate that dodder could indeed be used to transmit infectious clones. These results probed us to move the technology to woody hosts, were the dodder-based approach yielded significantly better results than direct agroinoculation, with BCRV efficiencies increasing from 9% to 73% and BYVaV from 0% to 46%.

Due to the unbalanced sample sizes across treatments, we chose to conduct a Fisher’s Exact Test to determine whether there was a significant association between the method used and the observed transmission. To assess the significance of the test results, a one-tailed p-value was calculated based on the output from Fisher’s test, allowing for a more detailed understanding of this association. Assays using *N. benthamiana* as the target host for BYVaV transfer revealed no significant differences between the dodder-based approach and the leaf agroinoculation method, indicating that both methods are equally effective. In contrast, for *Rubus* spp., statistical analyses provided strong evidence of an association between the transmission method employed and the observed transmission rates for both viruses. These findings suggest that the dodder method is significantly more efficient in facilitating virus transmission in *Rubus* and other woody hosts (blueberry, Hajizadeh, Atanda and Tzanetakis, unpublished data) when compared to agroinoculation.

It is known that dodder can transmit viruses (Bennett, 1956, Hosford, 1967, Li et al., 2022). Viruses that can invade the parenchymatous cells of the dodder haustorium can easily move to the host through perforations in the cell wall created during dodder-host interaction (Bennett, 1956). Moreover, the dodder haustorium not only establishes a direct connection between the vascular tissue of the host plant but also promotes the formation of plasmodesmata, linking the cytoplasm of dodder and host cells (Vaughn, 2003; Li et al., 2022). Li et al. (2022) further noted that the quantity and integrity of these plasmodesmata significantly impact the efficiency of mRNA transmission between plants (Li et al., 2022).

However, phloem-restricted viruses cannot utilize this invasion route. It is hypothesized that viral infection occurs due to a temporary reversal of food flow into the healthy host phloem, which allows the virus to enter the vascular system and initiate infection (Bennett, 1956; Hosford, 1967). The transmission of other phloem-restricted viruses using dodder, such as grapevine leafroll-associated virus-7, has been reported (Mikona & Jelkmann, 2010). We hypothesize that the higher transmission efficiency observed for the systemic BCRV, compared to the phloem-restricted BYVaV, is likely due to more accessible invasion routes.

This work presents an innovative approach for delivering infectious clones to woody hosts, utilizing dodder as an intermediary to transfer the virus vector from agro-inoculated plants to a target host. While dodder has been used in the past to facilitate virus transfer, this is the first report of its use in transmitting infectious clones. We successfully facilitated the transmission of infectious clones from two viruses to *Nicotiana benthamiana* and *Rubus* spp. showcasing the versatility of this method. This approach will facilitate the study of virus biology and host interactions across a broader range of plant species, contributing to developing more resilient crops.

## Authorship contribution statement

ASM and IET conceived and designed the experiments, ASM, MH and HA conducted experiments. ASM and IET analyzed the data. All participated in writing the paper and internal review. All authors have read and approved the final manuscript.

## Acknowledgements

We thank Dr. Richard Adams and Jenniffer Roa Lozano (University of Arkansas) for helpful discussion on the statistical analysis methodology. We also thank Dr. Dan Villamor for the thorough review and insightful suggestions on the manuscript.

## Funding

IET was supported by the United States National Institute of Food and Agriculture project ARK02850 and the Arkansas Agricultural Experimental Station.

## Declaration of generative AI and AI-assisted technologies in the writing process

During the preparation of this work the authors used ChatGPT v.4 to improve the language. No content was directly generated by the tool; rather, it was used to help enhance the overall clarity of the text. After using this tool, the authors reviewed and edited the content as needed and take full responsibility for the content of the publication.

---

**Supplementary Table 1.**
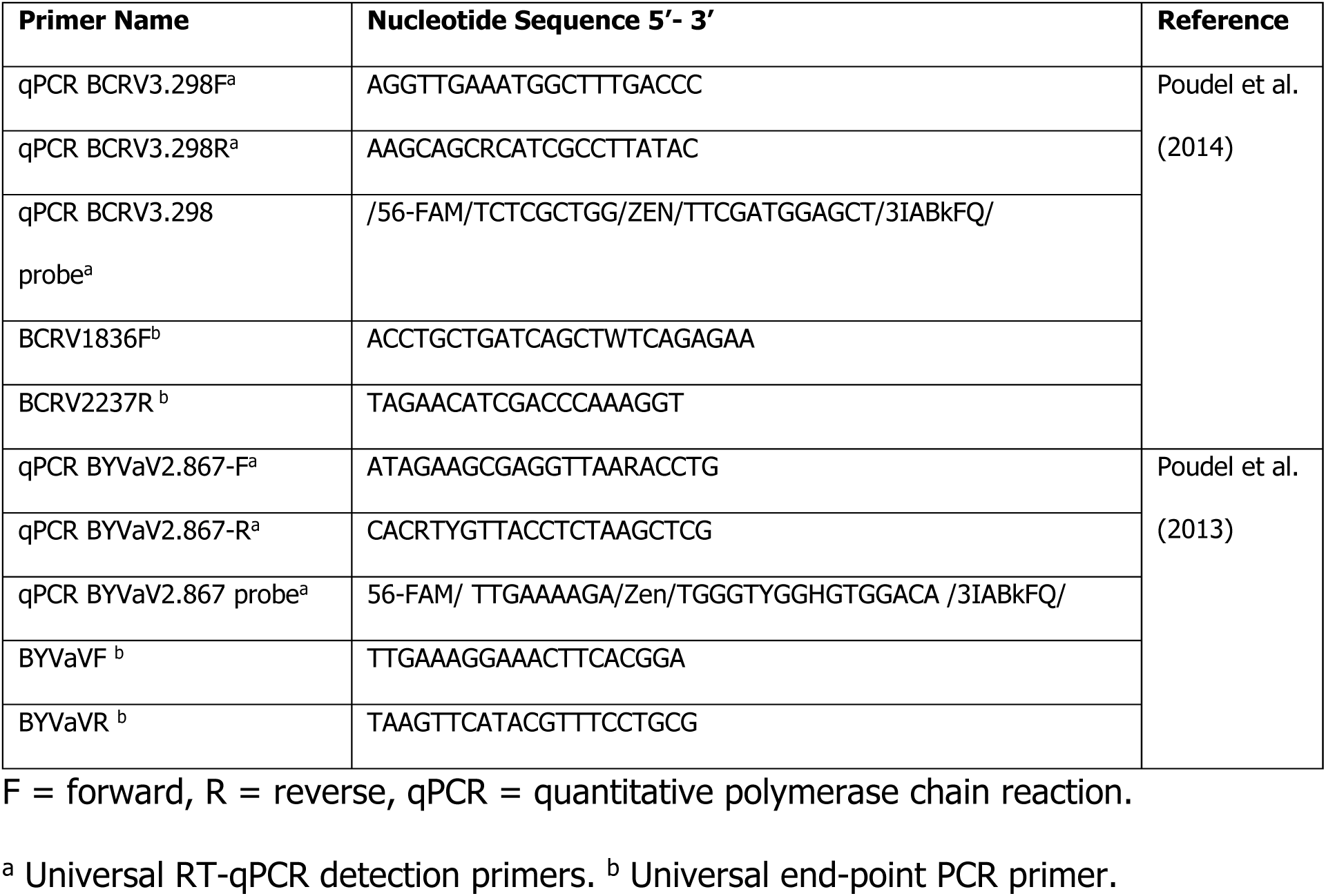
Detection primers for blackberry chlorotic ringspot virus (BCRV) and blackberry yellow vein associated virus (BYVaV).

## Notes

### Competing Interest Statement

The authors have declared no competing interest.

